# Diversity, Origin and Evolution of the ESCRT Systems

**DOI:** 10.1101/2024.02.06.579148

**Authors:** Kira S. Makarova, Victor Tobiasson, Yuri I. Wolf, Zhongyi Lu, Yang Liu, Siyu Zhang, Mart Krupovic, Meng Li, Eugene V Koonin

## Abstract

Endosomal Sorting Complexes Required for Transport (ESCRT) play key roles in protein sorting between membrane-bounded compartments of eukaryotic cells. Homologs of many ESCRT components are identifiable in various groups of archaea, especially in Asgardarchaeota, the archaeal phylum that is currently considered to include the closest relatives of eukaryotes, but not in bacteria. We performed a comprehensive search for ESCRT protein homologs in archaea and reconstructed ESCRT evolution using the phylogenetic tree of Vps4 ATPase (ESCRT IV) as a scaffold, using sensitive protein sequence analysis and comparison of structural models to identify previously unknown ESCRT proteins. Several distinct groups of ESCRT systems in archaea outside of Asgard were identified, including proteins structurally similar to ESCRT-I and ESCRT-II, and several other domains involved in protein sorting in eukaryotes, suggesting an early origin of these components. Additionally, distant homologs of CdvA proteins were identified in Thermoproteales which are likely components of the uncharacterized cell division system in these archaea. We propose an evolutionary scenario for the origin of eukaryotic and Asgard ESCRT complexes from ancestral building blocks, namely, the Vps4 ATPase, ESCRT-III components, wH (winged helix-turn-helix fold) and possibly also coiled-coil, and Vps28-like domains. The Last Archaeal Common Ancestor likely encompassed a complex ESCRT system that was involved in protein sorting. Subsequent evolution involved either simplification, as in the TACK superphylum, where ESCRT was co-opted for cell division, or complexification as in Asgardarchaeota. In Asgardarchaeota, the connection between ESCRT and the ubiquitin system that was previously considered a eukaryotic signature was already established.

**Importance:** All eukaryotic cells possess complex intracellular membrane organization. ESCRT (Endosomal Sorting Complexes Required for Transport) plays a central role in membrane remodeling which is essential for cellular functionality in eukaryotes. Recently, it has been shown that Asgard archaea, the archaeal phylum that includes the closest known relatives of eukaryotes, encode homologs of many components of the ESCRT systems. We employed protein sequence and structure comparisons to reconstruct the evolution of ESCRT systems in archaea and identified several previously unknown homologs of ESCRT subunits, some of which can be predicted to participate in cell division. The results of this reconstruction indicate that the Last Archaeal Common ancestor already encoded a complex ESCRT system that was involved in protein sorting. In Asgard archaea, ESCRT systems evolved towards greater complexity, and in particular, the connection between ESCRT and the ubiquitin system that was previously considered a eukaryotic signature was established.

## Introduction

Membranes are a fundamental staple of life. All cells are membrane-bounded, and membrane compartments, most likely, predated the emergence of the translation apparatus and even genetic information itself (1, 2). Genomes likely evolved within membrane-bounded protocells, and to ensure accurate segregation of replicating genomes into the daughter cells, a membrane-associated division apparatus had to evolve at the earliest stages of evolution. Accordingly, the molecular machinery involved in cell division could be expected to be conserved in all domains of life. However, this is far from being the case. Even if common structures, such as the division ring, are formed by most dividing cells, the proteins involved in this process differ (3–5). Even the most broadly conserved division machinery component, the FtsZ/tubulin GTPase involved in the division ring formation, is missing in several lineages of bacteria and archaea, and its function is taken over by other proteins (6–8). Most of the other protein components of the FtsZ-centered divisome, with a notable exception of SepF (9, 10), are not conserved but rather, are lineage-specific (3, 5, 11). Moreover, FtsZ paralogs can be repurposed to perform other functions related to membrane remodeling (12–14).

Extant living cells not only employ diverse proteins for division, but form vesicles and intracellular compartments that require specific membrane remodeling systems, often distinct from the cell division apparatus (15). Eukaryotic cells possess a highly complex cell division machinery and multiple endomembrane organelles, such as mitochondria, endoplasmic reticulum, Golgi complex and others. The traffic of molecules between these compartments in eukaryotic cells is enabled by several sorting systems (16). One of such molecular machines is ESCRT (Endosomal Sorting Complexes Required for Transport), which mediates a variety of membrane scission events, including a key role in cytokinesis in at least some plant and animal cells (17, 18). The composition, structure and function of ECSRT complexes in eukaryotes have been studied in detail (17, 19–21). The core of this system consists of four interacting, multisubunit complexes that are conserved in most eukaryotes (22): ESCRT-I, -II, -III, -IV and the less deeply conserved ESCRT-0 (Figures 1 and 2). The ESCRT complex is linked to other protein trafficking systems, such as adaptins and clathrins (Figure 1).

**Figure 1.**
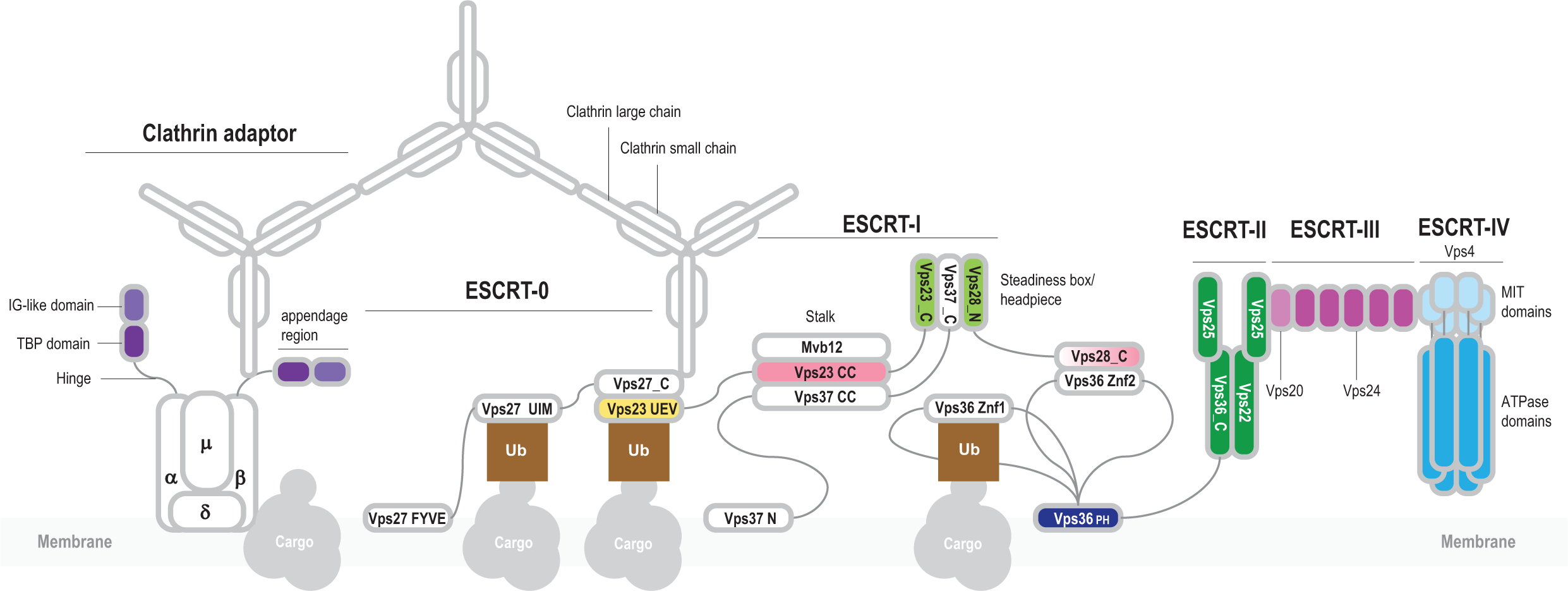
General organization of the eukaryotic ESCRT system and its connection with other protein sorting complexes. The ESCRT machinery was thoroughly characterized in yeast and mammals (17, 19–21). The core of the system consists of four conserved, multisubunit complexes: ESCRT-I, -II, - III, -IV. ESCRT-0 subunits are not generally conserved and apparently differ among the eukaryotic phyla. ESCRT-0 subunits interact with protein cargo and recruit ESCRT-I complex. The best characterized subunit of ESCRT-0 is Vps27 which is a hub of protein-protein interactions (19). Vps27/HRS is a multidomain protein containing a VHS (Vps27/HRS/STAM) super-helix domain, a FYVE zinc finger, several coiled-coil regions and a C-terminal P[ST]xP motif. Vps27 interacts with the Ubiquitin E2 variant (UEV) domain of Vps23 (ESCRT-I) via the P[ST]xP motif, and also with clathrin, ubiquitinated cargo and lipids (19, 97, 98). ESCRT-I consists of four subunits: Vps28, Vps23, Vps37 and Mvb12 (Figure 2). Three subunits, Vps23, Vps37 and Mvb12, form a stalk through the interaction between three long helices forming a coiled-coil structure. Vps23, Vps28, Vps37 form a headpiece through the interaction between homologous helical hairpins known as the Steadiness Box (SB) (99). The C-terminal helical bundle domain of Vps28 interacts with ESCRT-II Vps36 via a Zn finger (Znf2) inserted into the split PH domain. ESCRT-II also consists of four subunits: Vps22, Vps36 and two copies of Vps25 (100). These proteins are paralogs that share a common region consisting of a pair of wH domains. In addition to the wH domains, Vps36 is fused to a PH domain and two zinc finger domains (the latter are absent in mammalian orthologs). Vps22 and Vps36 contain an additional helical domain upstream of the proximal wH domain, which likely promotes their interaction (Figure 2). The N-terminal domain of VPS25 binds the C-terminal domain of VPS22, and the other VPS25 subunit contacts both VPS36 and VPS22, forming an asymmetric Y-shaped structure (100). Vps25 recruits the initial Vps20 subunit of ESCRT-III complex (100). ESCRT-III consists of several paralogous subunits, which belong to two groups, Vps2-like (MIM-1 [MIT-interacting motif] containing) and Vps20-like (MIM-2 containing), respectively ^16,29^ (Figure 2). Vps2 is targeted to the membrane where it nucleates Vps20 polymerization^16^. Depolymerization of ESCRT-III is regulated by the Vps4^16^. Vps4 contains a diagnostic N-terminal three-helix bundle, the microtubule interacting and trafficking (MIT) domain which interacts with the C-terminal MIM-2 in ESCRT-III subunits (101). The ESCRT machinery is required for endosomal trafficking and is connected with other complexes involved in this process including clathrin, a large complex consisting of a light and a heavy subunits forming a triskelion (102). In eukaryotes, clathrin forms a cage-like scaffold around a vesicle and recruits Vps27. Clathrin is associated with clathrin adaptor complexes (AP) which mediate clathrin-dependent protein trafficking to and from endosomes (44, 50). AP complexes differ minimally in subunit composition. AP2, for example, consists of four subunits: alpha, beta, gamma and mu (44, 50). Alpha and beta subunits are paralogs that contain an N-terminal HEAT repeats domain and a C-terminal appendage domain connected through an unstructured hinge interacting with clathrin. The appendage domain consists of two distinct subdomains, the proximal immunoglobulin-like (IG) beta sandwich fold, and the distal TBP (TATA-box binding protein or helix-grip) fold (103). Appendage domains of alpha and beta subunits can recruit multiple additional proteins, whereas the N-terminal domains of alpha and beta subunits and the mu subunit interact with the membrane protein cargo (66, 67).

**Figure 2.**
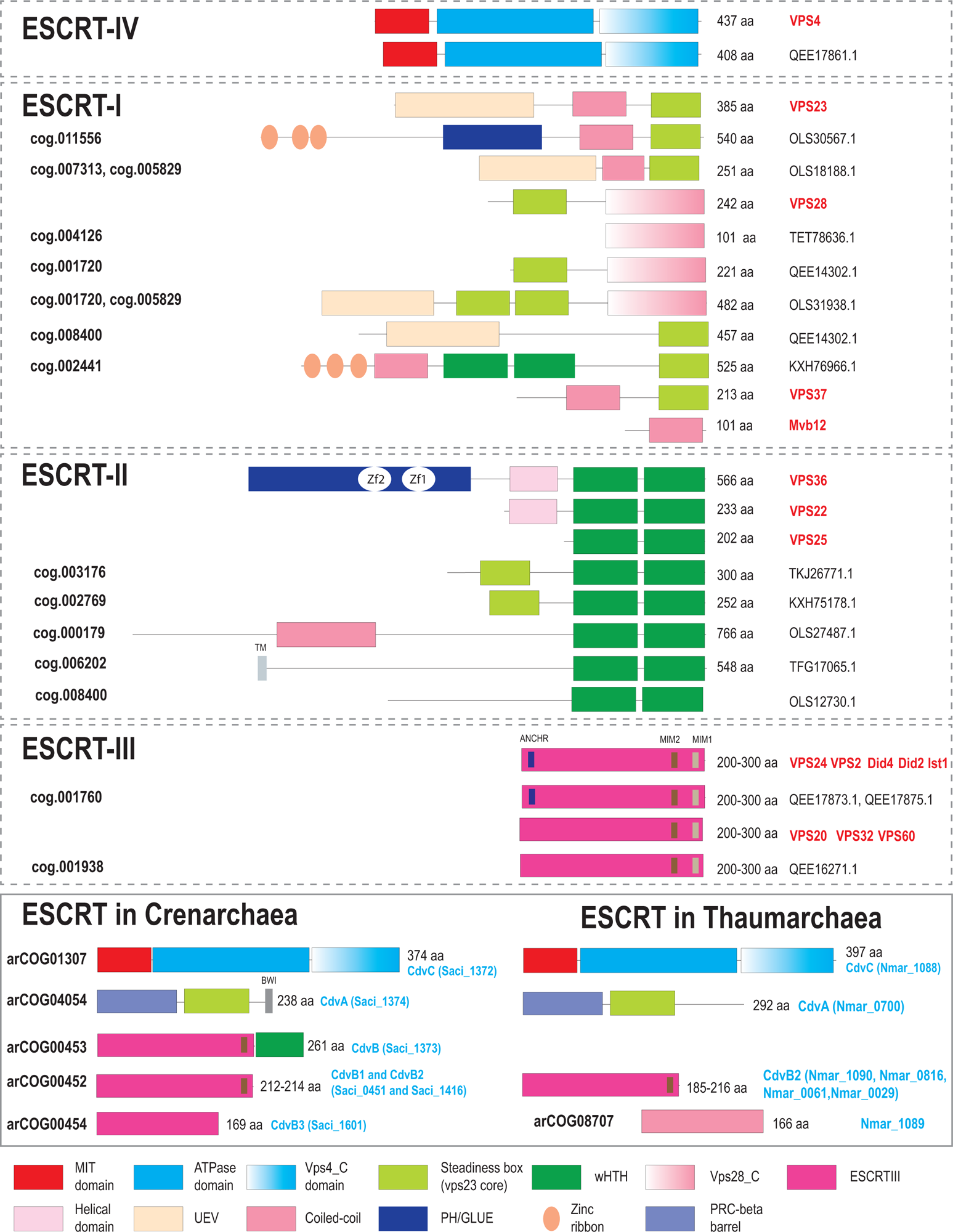
Domain organization and key motifs of ESCRT protein in eukaryotes, Asgard and TACK archaea. Protein domains are shown as colored shapes according to the color code given beneath the schemes. Proteins are drawn roughly proportional to their size in amino acids which is indicated in the right. Domain boundaries for *Saccharomyces cerevisiae* are based on previously published structures and analyses (19, 55, 98). For archaeal proteins, domain boundaries are based either on previous analyses (22, 35, 53, 54) or on sequence analysis performed during this work as described under Methods. *S. cerevisiae* gene names are highlighted in red on the right. For Asgard and TACK archaea, protein size and accession are indicated on the right. asCOG numbers (27) are indicated for the Asgard proteins and arCOG numbers (34) are indicated for the TACK proteins on the left. Signature sequence motifs are shown by small rectangles, and short names for the motifs are provided above the protein schematics (once for proteins with the same domain organization). Abbreviations: Zf, Zinc finger; TM, Transmembrane segment; UEV, Ubiquitin E2 Variant; MIM, MIT Interaction Motif; BWI, Broken Winged-Helix Interaction Site. Not all proteins from cog.003176 are fused to SB, but most contain additional N-terminal subdomains distinct from those in eukaryotic homologs.

For a long time after their discovery, protein components of ESCRT complexes were thought to be eukaryote-specific (23). However, in 2008, Lindås et al discovered a cell division (Cdv) machinery in Crenarchaeota that was distinct from the previously characterized FtsZ based system present in most bacteria and archaea. CdvB and CdvC proteins were found to be homologous to eukaryotic ESCRT-III and IV (Vps4) subunits, respectively, whereas no homologs were identified for CdvA (8). Subsequently, ESCRT-III components and the Vps4 ATPase, another essential ESCRT component, were identified in a wider range of archaea (24). However, the distribution of these proteins is patchy, and they were found to be conserved only in Sulfolobales, Desulfurococcales and Thaumarchaea (24). Nevertheless, these findings suggested deep archaeal roots for ESCRT-III and Vps4. More recently, structural and functional similarities were detected between polymers of ubiquitous bacterial membrane remodeling proteins Vipp1 and PspA, and those of eukaryotic and archaeal ESCRT-III subunits, suggesting that ancestors of ESCRT-III were present in the Last Universal Cellular Ancestor (LUCA) (25). However, the origin of ESCRT-I and II remained enigmatic until the discovery of Asgardarchaeota (Asgard for short), the phylum that includes the closest known archaeal relatives of eukaryotes which, in most phylogenetic trees of conserved informational proteins, are associated with the Asgard class Heimdallarchaeia (26–28). Strikingly, homologs of the subunits of ESCRT-I, -II, -III and -IV complexes along with a complete ubiquitin machinery that plays a key role in protein fate determination in eukaryotes were identified in Asgard archaea (27–30).

A closer examination of the components of the ESCRT complex revealed remarkable diversity of domain architectures in Asgard archaea (27). In view of this diversity and the relatively low sequence conservation among ESCRT components, we re-analyzed archaeal ESCRT systems, in particular, taking advantage of AlphaFold2 (31) (AF2) structure predictions for analysis beyond the sequence level. This analysis resulted in the identification of several distinct groups of ESCRT systems in archaea outside of Asgard, including proteins structurally similar to ESCRT-I and ESCRT-II components, and several other domains involved in protein sorting in eukaryotes, suggesting an early origin of these components. Additionally, we identified distant homologs of CdvA in Thermoproteales which are likely to be components of the elusive cell division system in this group of archaea. We propose an evolutionary scenario for the origin of eukaryotic and Asgard ESCRT complexes from ancestral building blocks, namely, the Vps4 ATPase, ESCRT-III components, wH (winged helix-turn-helix fold) and possibly also coiled-coil, and Vps28-like subdomains.

## Results and Discussion

### Phylogeny of Vps4 ATPase and distribution of ESCRT systems across archaea

Vps4 family ATPase is present in all ESCRT systems and is conserved at the sequence level, making it suitable for phylogenetic reconstructions and for analyzing ESCRT evolution. We identified Vps4 (to the exclusion of other AAA+ ATPases) in archaea and eukaryotes based on two criteria: 1) presence of the Microtubule Interacting and Trafficking (MIT) domain, known to be essential for interaction with ESCRT-III, or 2) presence of genes encoding ESCRT components in the respective archaeal genome neighborhoods. Using these criteria, we compiled a representative Vps4 set that included 97 proteins from Asgard archaea, 152 proteins from other archaea (arCOG01307), and 71 proteins from selected eukaryotes (Supplementary Table 1, Supplementary File 1). A phylogenetic tree of Vps4 was built from the multiple alignment of these sequences (Figure 3A). The tree was rooted using the related but distinct Cdc48 ATPases as an outgroup (see Methods and Supplementary Figure 1).

**Figure 3.**
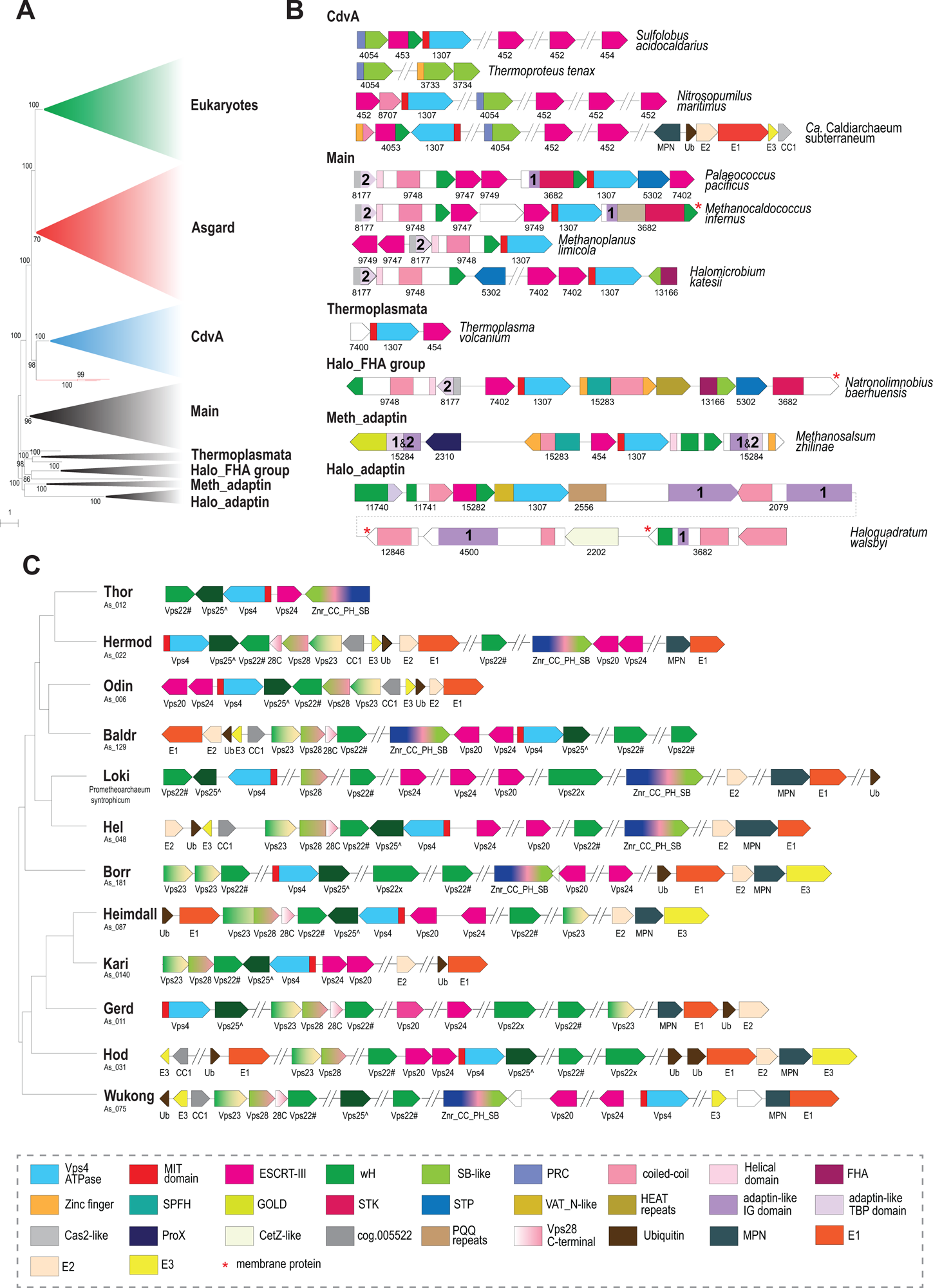
Phylogeny of Vps4 family ATPase and genomic neighborhoods of ESCRT systems in archaea. **A. Schematic representation of the phylogenetic tree of Vps4 protein family.** The phylogenetic tree was built using IQ-Tree as described in Material and Methods. Major branches were collapsed and are designated on the right. Bootstrap values > 70% calculated by IQ-Tree are shown. The complete tree in Newick format is available in the Supplementary File 1. **B. Organization of selected ESCRT loci for each major branch of archaea (except Asgard).** Genes are shown as arrows roughly proportional to gene size. Domains are indicated within the arrows for multidomain proteins, not to scale. Homologous genes and domains (or generic structural elements such as coiled-coil or helical domains) are color coded according to the code at the bottom of the figure. ArCOG numbers are indicated below the arrows. Numbers inside the arrows indicate IG domain (1) and TBP domain (2) in the adaptin appendage homology regions. **C. Components of ESCRT systems in selected genomes of 12 major Asgard lineages.** The ESCRT related genes are shown for selected genomes of the 12 major lineages of Asgradarchaeota(27). The tree schematically shows the relationships among the lineages according to previously published phylogenetic analysis(27). Designations are the same as in B, except for multidomain proteins in which domains are not shown, but are explained in the color code schematics in the bottom of the figure. Protein names are indicated below the arrows. Designations: Vps22# - Asgard specific version of Vps22, cog.002769; Vps25^ - Asgard specific version of Vps25, cog.002441; Znr_CC_PH_SB – Asgard specific protein family of cog.011556 (see Figure 2). Abbreviations: wH - winged helix domain, Znr - zinc ribbon, SB - Steadiness box, FHA - forkhead-associated domain, STK - serine/threonine protein kinase, STP - serine/threonine protein phosphatase, UEV - Ubiquitin E2 variant; Ub – ubiquitin, CC-coiled-coil; PH-plextrin homology; SB – steadiness box; MIT - microtubule interacting and transport; HEPN - higher eukaryotes and prokaryotes nucleotide-binding domain; GOLD - Golgi dynamics domain; SPFH - stomatin, prohibitin, flotillin, and HflK; VAT_N – N-terminal domain of valosin-containing protein-like ATPase of *Thermoplasma acidophilum*; TBP-TATA-binding protein. Abbreviations for repetitive domains: HEAT - Huntington, Elongation Factor 3, PR65/A, TOR; PQQ – pyrrolo-quinoline quinone, PEGA – containing PEGA sequence motif. Abbreviations for ubiquitination pathway: E1 - ubiquitin-activating enzyme; E2 - ubiquitin-conjugating enzyme; E3 - ubiquitin ligases; MPN - Mpr1/Pad1 N-terminal domain, deubiquitinating enzyme.

Eukaryotic and Asgard Vps4s were found to be monophyletic except for three diverged paralogs from Hodarchaeota that group outside of the main Asgard clade (Figure 3A, Supplementary File 1). Considering that the same Hodarchaeotal genomes encoded Vps4 variants that group within the main Asgard clade and that these three outliers are not associated with any other ESCRT system genes, they likely represent a fast evolving, subfunctionalized subfamily. The Eukaryotic-Asgard clade is a sister group to a large clade corresponding to Vps4s from the TACK (Thaumarchaeota [now Nitrososphaeria], Aigarchaeota, Crenarchaeota [both Thermoproteota], and Korarchaeota [now Korarchaeia]) superphylum. In these organisms, ESCRT systems have been shown to participate in cell division (4, 8, 32) and extracellular vesicle biogenesis (33). We denoted this clade CdvA, after the unique protein diagnostic of these systems. Most of the remaining, largely euryarchaeal, Vps4s form 5 clades. The largest and most taxonomically diverse clade was denoted “Main”. As noted previously (24), several Thermococcales species encode a solo Vps4, but not other components of the system (Supplementary Figure 1, Supplementary Table 2). These standalone Vps4 proteins belong to a separate clade on a long branch suggestive of subfunctionalization.

The tree included also a Thermoplasmata-specific clade, and three clades that we denoted by their distinct features and the respective taxa: Halo_FHA (Halobacteriales, FHA ([forkhead-associated] domain containing); Meth_adaptin (Methanobacteriales, adaptin appendage domain containing) and Halo_adaptin (Halobacteriales, adaptin-like IG [immunoglobulin] domain containing) (Figure 3A).

The Main clade included representatives from Thermococcales, Methanococcales, Methanomicrobiales and Halobacteriales orders and several other archaea (Supplementary File 1). Besides Vps4 and ESCRT-III, all systems in this clade encode two proteins that belong to arCOGs (34) 09748 and 08177 (Supplementary Table 2). arCOG09748 proteins contain a C-terminal wH domain with sequence and structural similarity (Z-score=6.4) to the wH domain at the C-terminus of CdvB (35) and structural similarity (Z-score=6.7) to the C-terminal domain of Vps22 (ESCRT-II complex) (Figure 4A, Supplementary Figure 2, Supplementary Table 3). AF2 modeling for arCOG09748 protein from *Palaeococcus pacificus* revealed an N-terminal helical domain, a coiled-coil domain, a globular central domain (no structural similarity to any known domains) and a wH C-terminal domain (Supplementary Figure 2). The N-terminal domain of arCOG08177 proteins showed surprising similarity to an apparently inactivated Cas2 nuclease, the CRISPR-Cas system component that forms a complex with the Cas1 integrase and is involved in spacer acquisition (36) (Supplementary Figure 2, Supplementary Table 3). Like Cas2 in the CRISPR adaptation complex, the N-terminal domain of arCOG08177 proteins could mediate dimerization of arCOG08177 proteins (36). The second domain of arCOG08177 corresponds to the archaea-specific domain DUF3568 and adopts the TBP (TATA Binding Protein or helix-grip) fold shared with the C-terminal domain of the adaptin appendage region (Figure 3B and 4) (37). In many genomes from the Main clade, the ESCRT locus additionally contained genes encoding serine/threonine protein kinase (STK) and protein phosphatase (STP) (Figure 3B, Supplementary Table 2). Typically, these STKs are membrane associated. Most STK consist of several domains: a transmembrane segment at the N-terminus, immunoglobulin-like repeats (found in the appendage region of adaptins) domain and a C-terminal small helical domain for which HHpred search revealed similarity with a distinct variant of the wH fold, PCI (for Proteasome, COP9, Initiation factor 3) domain (Figure 4B, Supplementary Figure 2, Supplementary Table 3). The PCI domain is responsible for oligomerization of these proteins in respective complexes (38, 39). Thus, this domain in STK could be involved in either oligomerization or in interaction with wH domain of arCOG09748 proteins.

**Figure 4.**
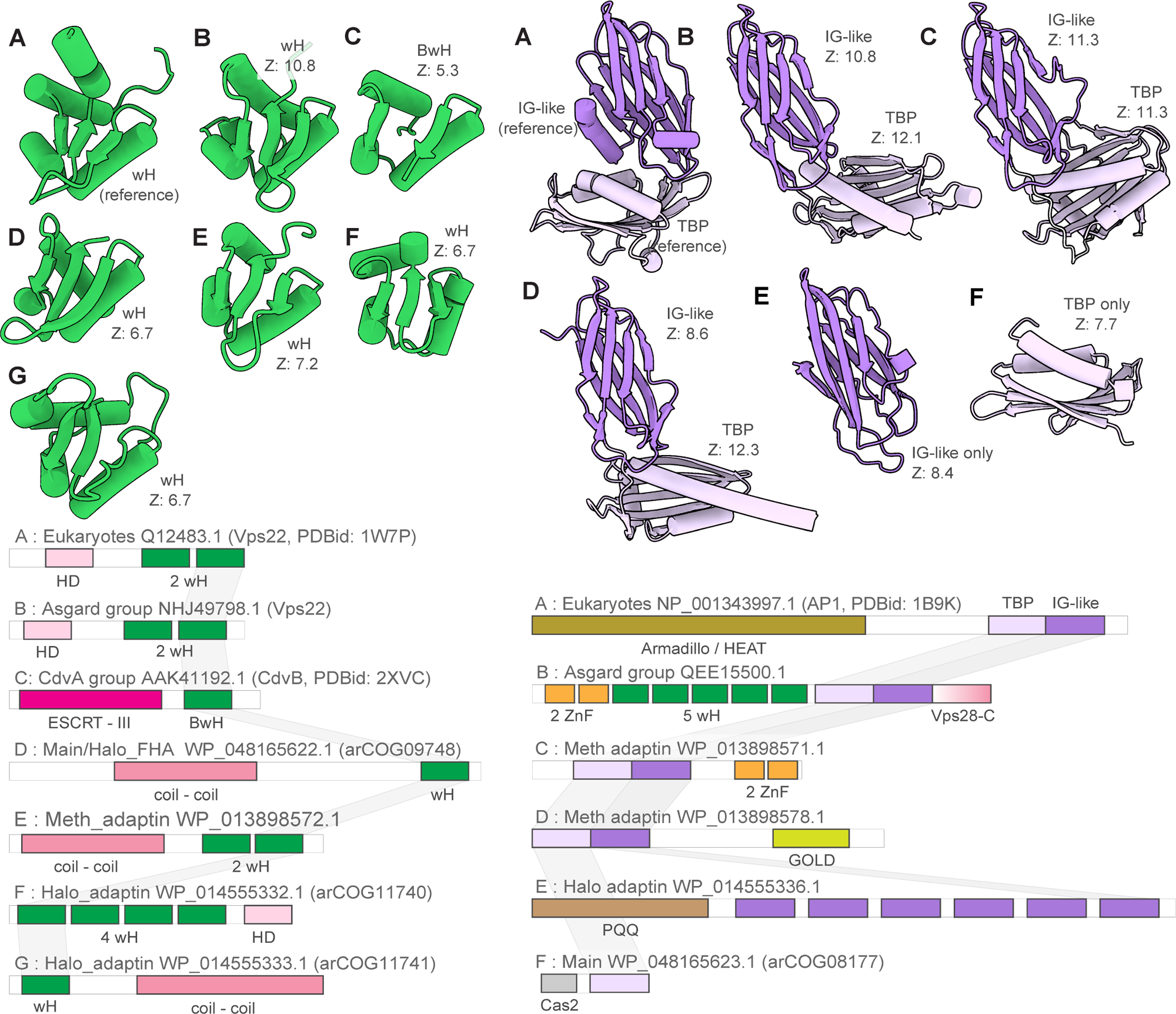
Winged helix and adaptin appendage domain homologs in eukaryotic and archaeal ESCRT systems. Left, comparison of winged helix domains (wH). Right, adaptin appendage-like regions. Multidomain proteins were trimmed to show only domains homologous to wH and adaptin appendage-like domain of Vps22 and alpha-adaptin appendage domain. Complete proteins are shown in Supplementary Figures 2, 7 and 8. Complete domain organizations of the respective proteins are shown underneath the structures. Letters denote the correspondence between structures and domain architectures. DALI Z-scores between the indicated domains and wH domain of Vps22 or IG-like and TBP domain of alpha-adaptin appendage domain are indicated. Abbreviations: IG, Immunoglobulin; wH, winged helix domain; BwH, Broken winged Helix domain; TBP, TATA binding protein

The Thermoplasmata clade included minimal ESCRT systems that consist of genes encoding Vps4, ESCRT-III and a beta-barrel protein from arCOG07400 (Figure 3B, Supplementary Table 1 and 2). Several Thermoplasmata species lack FtsZ but encode complete or partial ESCRT systems that are likely involved in cell division (Figure 5).

**Figure 5.**
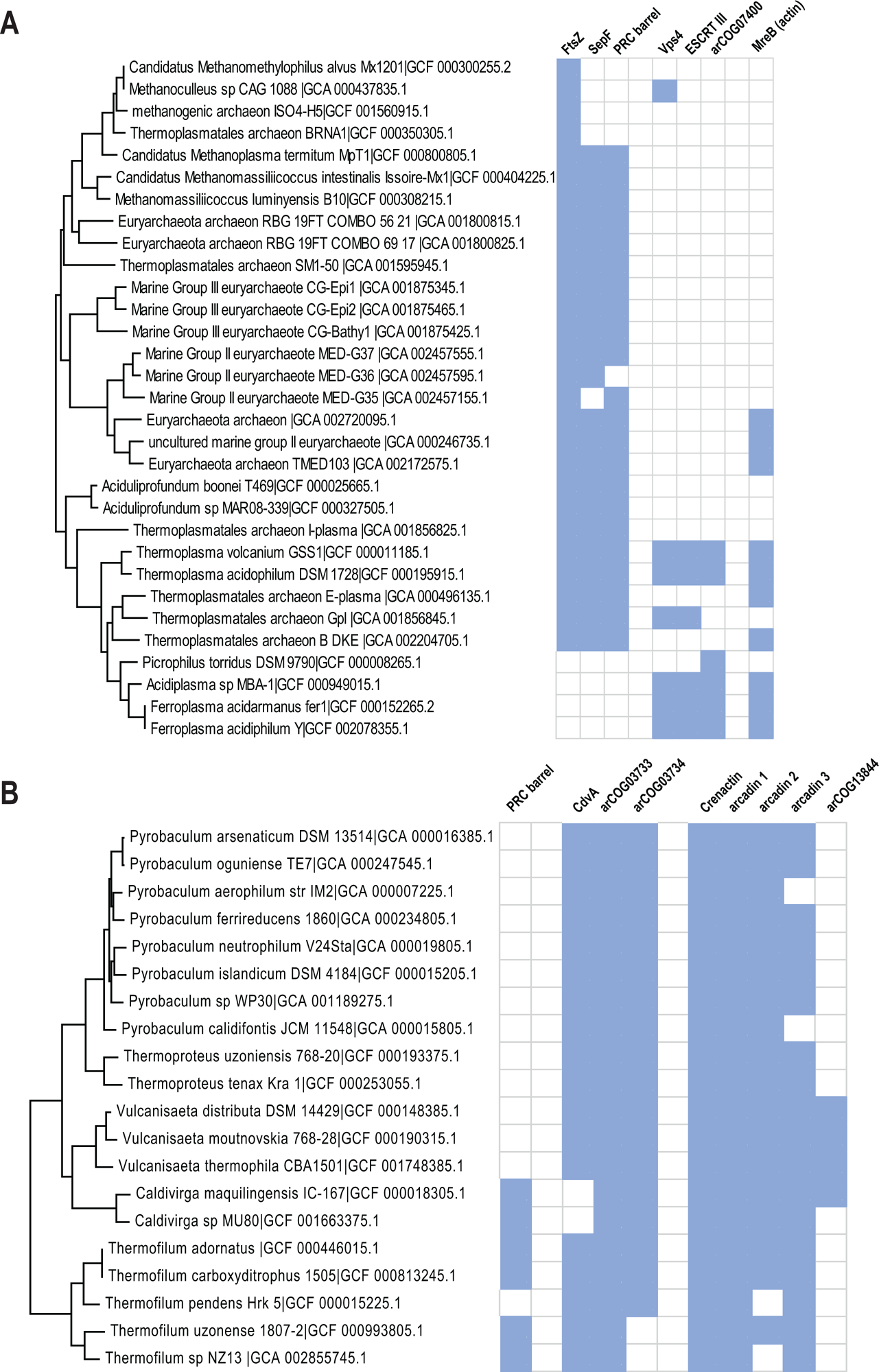
Phyletic patterns for putative cell division components of Thermoproteales and Thermoplasmata. The presence/absence patterns of cell division components for **A)** Thermoplasmata and **B)** Thermoproteales are shown. The patterns are overlayed on the respective species subtree. The tree built for all 524 arCOGs genomes using IQ-tree program (91) and concatenated alignment of ribosomal proteins. Blue indicates presence, and white indicates absence.

A distinct FHA domain family (arCOG13166) is associated with Vps4 and ESCRT-III in several haloarchaeal genomes comprising the signature of the Halo_ FHA clade (Figure 3B, Supplementary Figure 1, Supplementary Table 2). The FHA domain is involved in phosphorylation-dependent protein oligomerization and/or interaction with phosphorylated proteins (40, 41). The C-terminal domain of arCOG13166 proteins showed limited similarity to CdvA (Figure 3B, Supplementary Figure 4 and 5, Supplementary Table 3). Proteins of arCOGs 09748 and 08177, STKs and STPs are also encoded in some of these Halo_FHA loci (Figure 3B, Supplementary Table 2).

Meth_adaptin neighborhoods encode two proteins similar to the adaptin appendage region (both IG-like and TBP domains) (Figure 3B and 4B, Supplementary Figure 6, Supplementary Tables 2 and 3). One of the adaptin appendage region containing proteins (eg. WP_013898571.1) also contains a Zn-ribbon domain and an additional uncharacterized N-terminal domain (Supplementary Figure 6). In one of these proteins (eg. WP_013898578.1), the appendage region is additionally fused to a GOLD (Golgi dynamics) domain(42) via a transmembrane linker (Supplementary Figure 6). GOLD domains connect membrane cargo proteins with COPI (an adaptin homolog) vesicles (43, 44). Another component of the Meth_adaptin systems belongs to the SPFH (stomatin, prohibitin, flotillin, and HflK/C) family (Figure 3B). In eukaryotes, SPFH proteins interact with the ESCRT-0 component Vps27 which presents ubiquitinated proteins to the ESCRT machinery (45). SPFH-family protein additionally contains coiled-coil and zinc ribbon-domain (Supplementary Figure 6). Interestingly, bacterial SPFH-family protein YdjI, is encoded in the some PspA loci (ESCRT III family) and is required to localize PspA to the membrane (46), suggesting that the functional connection between ESCRT III and SPFH proteins could have been established very early in evolution. Recently, a structure of a distant bacterial SPTH homolog has been solved and it has been shown that it forms 24-mer with coiled-coil domains interacting and forming a “lid” of an enclosed compartment on a membrane (47). This protein might play a similar role in ESCRT systems. Finally, Meth_adaptin loci included a gene encoding ProX, a substrate-binding protein of the osmoregulatory ABC-type glycine betaine/choline/L-proline transport system (48) (Figure 3B, Supplementary Table 2). The ProX-like protein likely binds a substrate which is sorted into vesicles formed by other genes of this ESCRT system.

Despite grouping with the Meth_adaptin clade in the tree, the architecture of Halo_adaptin loci is quite different (Figure 3B). In these systems, the Vps4-like ATPase (eg. WP_014555335.1) lacks an identifiable MIT domain, and instead contains an N-terminal domain related to the N-terminal domain of the Cdc48-like ATPase VAT (valosin-containing protein of *Thermoplasma acidophilum*) (Supplementary Table 3). Upstream of the ATPase, there is a gene encoding a protein with weak sequence similarity to PspA, an ESCRT-III superfamily protein (Supplementary Table 4, Supplementary Figure 7). Given that in most other archaeal ESCRT systems, at least one ESCRT-III gene is located upstream of Vps4, this protein is likely to be a derived version of ESCRT-III (Figure 3B; Supplementary Figure 7). These neighborhoods also encompass genes encoding two wH containing proteins: arCOG11740 and arCOG11741. As in many other systems described above, a wH protein (arCOG11741) is encoded upstream of the ESCRT III-like protein suggesting that these proteins interact (Figure 3B, Supplementary Table 2). The wH domain in arCOG11741 proteins is fused to a coiled-coil domain. The wH containing arCOG11740 proteins are encoded by the proximal gene in the loci (Supplementary Table 2). Like eukaryotic Vps36, arCOG11741 proteins contain duplicated wH domains and an N-terminal alpha-helical domain (Figure 3B and 4A, Supplementary Figure 7, and 3). Similarly to the Meth_adaptin systems, these neighborhoods encode two distinct proteins with sequence and structural similarity to the IG-like domain of the adaptin appendage region (Supplementary Figure 7, Supplementary Table 3). Both “adaptin-like” proteins in Halo_adaptin system contain several IG-like domains (Supplementary Figure 7). The N-terminal domains of both “adaptin-like” proteins adopt a WD40/PQQ-like beta propeller fold (Supplementary Figure 7). Notably, a WD40-like domain is present in two homologous COPI alpha and beta-prime subunits, which mediate interaction with specific protein cargo (49) and in the heavy subunit of clathrin (50). Halo_adaptin systems are the most complex and additionally include two more IG-like domains containing proteins, a membrane protein and a protein related to CetZ (FtsZ/tubulin family) that is involved in cell shape control in *Haloferax volcanii* (14) (Figure 3B, Supplementary Table 2). Phylogenetic analysis showed that the CetZ-like protein from *Haloquadratum walsbyi* groups outside the main CetZ branch suggesting a different function (14). Here we link this function to the Halo_adaptin system likely involved in protein sorting.

Most members of the TACK superphylum possess a minimal ESCRT system consisting of CdvA, CdvB (ESCRT-III) and CdvC (Vps4) (Figure 3B, Supplementary Table 1). Recently, CdvA orthologs have been identified in almost all Thermoproteales (51) where they were previously thought to be missing. In addition to CdvA orthologs found in most Thermoproteales, we report here two more CdvA-related protein families specific for this lineage, arCOG03733 and arCOG03734 (Figure 5B, Supplementary Figure 4, Supplementary Table 3). These two paralogs are encoded next to each other in most of the Thermoproteales genomes (Supplementary Table 2). Proteins of arCOG03733 additionally contain a Zn ribbon N-terminal domain connected to CdvA-like domain through a long coil-coiled like region, whereas proteins of arCOG03734 are closely similar but lack the Zn ribbon. The domain architectures of these proteins, along with CdvA, are similar to that of eukaryotic Vps23 and Vps37 components of ESCRT-I (Figure 2). Furthermore, HHpred search revealed sequence similarity between the helical hairpin of the Steadiness Box (SB) and CdvA (Supplementary Table 3, Supplementary Figure 4), supporting our previous observations (52). Recent examination of CdvB protein sequences suggested that most of them harbor a C-terminal MIT interaction motif 2 (MIM2) that is conserved in Vps20/32/60 subunit class (53) (Figure 2). Most of the TACK genomes encode additional ESCRT III family proteins that, however, lack the wH extension as noticed previously (54). Furthermore, in thaumarchaea, none of the CdvB family proteins contain fused wH domains (54). Another distinctive feature of this system in thaumarchaea is the additional component (arCOG08707), a coiled-coil protein, encoded upstream of the *cdvC/vps4* gene (Figure 3B). arCOG08707 proteins are not similar to any known proteins, but structural models shows that they possess a helical hairpin which is present in both ESCRT III and CdvA (SB) (Supplementary Figure 4). Bathyarchaea encode additional CdvA-like proteins (eg. KPV61682.1) which are fused to an unstructured glutamate rich sequence and are encoded outside of the *cdv* locus.

### Extensive diversification of ESCRT in Asgard archaea

The diversity of ESCRT systems in Asgardarchaeota far exceeds that in any other archaea. Most of the Asgard lineages encode multiple eukaryotic-like ESCRT components as well as proteins involved in ubiquitination, suggesting interaction between the two systems, both of which are prominent in eukaryotes (22, 27, 28). In agreement with previous observations (22, 25, 27, 28, 54), we identified orthologs of the ESCRT III Vps24 and Vps20 families (Figure 2 and 6A). Compared to eukaryotes (19), Asgard archaea have fewer paralogs in both families.

**Figure 6.**
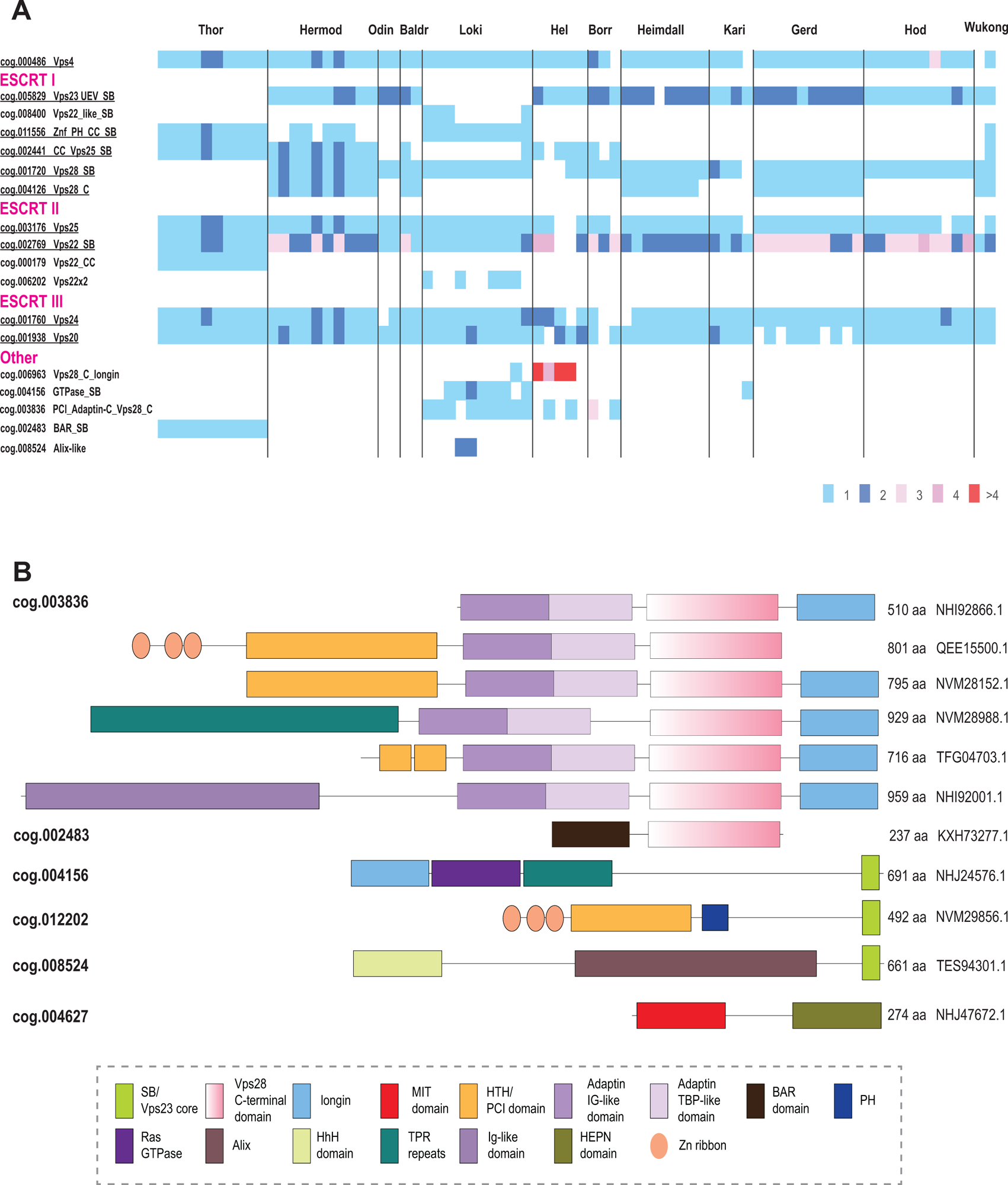
Phyletic patterns of ESCRT components and unique multidomain proteins in Asgardarchaeota. **A. Number of proteins in ESCRT asCOGs for 78 Agard phylum genomes.** Major Asgard lineages are indicated above. asCOG numbers and a short description or gene or protein name are indicated on the left; the data were retrieved from asCOGs ^23^ with minimal corrections. The numbers of proteins are color coded as shown in the bottom right. Potentially ancestral (most widespread) asCOG are highlighted in green. **B. Domain architectures of Asgard proteins containing one or more domain related to ESCRT system.** asCOG numbers are indicated on the left. Protein sizes and accession are indicated on the right. Homologous domains are shown by the same shape and color according to the color code provided in the bottom. Abbreviations are the same as in Figure 3. Additional abbreviations: BAR - BIN, amphiphysin and Rvs161 and Rvs167 (yeast proteins) domain; TPR, tetratricopeptide repeats.

The ESCRT-II proteins show considerable diversity in both domain organization and number of paralogs (Figure 2 and 5A). Unlike in eukaryotes, Vps22 family proteins (Asgard cog.002769) are fused to an SB, whereas Vps25 proteins (cog.003176) contain additional N-terminal subdomains some of which are also similar to SB (Figure 2, Supplementary Table 3).

Another frequent domain arrangement unique for Asgards (cog.002441) also includes two wH domains along with an SB and three Zn ribbons (Figure 2). Among the ESCRT-I components, Asgard archaea encode orthologs of both Vps23 and Vps28 (Figure 2, Figure 3C). In some Asgard genomes, the C-terminal domain of Vps28 is encoded separately, in addition to a full *vps28* gene (Figure 3C). Furthermore, we identified a protein with a distinct domain organization, consisting of three Zn ribbon domains, a PH domain, a coiled-coil region and an SB box (Znr_PH_CC_SB), that is absent in eukaryotes, but is found in several Asgard lineages and is likely to be ancestral in Asgardarchaeota (Figure 2 and 3C). Given the presence of SB, the distinct structural module of eukaryotic ESCRT-I, Znr_PH_CC_SB is a likely ESCRT-I component in Asgard archaea. In yeast, the PH domain of Vps36 interacts with phosphatidylinositol-3-phosphate, whereas one of the two Zn finger domains binds to ubiquitinated cargo and the other one interacts with ESCRT-I via the C-terminal domain of Vps28 (Figure 1) (55). By analogy, the PH domain of Znr_PH_CC_SB is expected to be membrane associated.

Typically, *vps25*-like genes are codirected (and likely co-translated) with *vps4* and ESCRT III, whereas *vps22*-like genes are co-directed with *vps23*-like and *vps28*-like genes as noted previously (22) (Figure 3C). This arrangement most likely reflects the order of assembly of these components. Unlike in eukaryotes, Asgard Vps28-like SB domain and Vps28_C terminal domain are encoded by two separate genes (Figure 3C). In eukaryotes, the Vps28 C-terminal domain interacts with ESCRT II complex subunits (56, 57), suggesting a similar interaction takes place in Asgard archaea.

Lokiarchaeia and Helarchaeia genomes harbor many different families and different domain organizations of ESCRT system proteins, whereas Hermodarchaeia, Hodarchaeia and Wukongarchaeia have none (Figure 6A). Such diversity of ESCRT systems implies their functional specialization in different groups of Asgard archaea.

In eukaryotes, ESCRT-0 components are highly diverse and differ between fungi and mammals, with only a few genes shared (19). Orthologs of the most highly conserved Vps27 or Bro1/Alix (apoptosis-linked gene-2 interacting protein X) ESCRT-0 proteins were not identifiable in Asgard archaea. Conversely, many multidomain proteins containing domains related to ESCRT core components were detected in Asgard, but not in eukaryotes (27). Most of these proteins contain either the Vps28 C-terminal domain (Vps28-C) or the SB (Figure 6B). The Vps28 C-terminal domain is often present along with the adaptin appendage region and a longin domain. Assuming that Vps28-C interacts with ESCRT-II, these proteins could associate with ESCRT-II without the involvement of any core ESCRT-I subunits. The presence of longin suggests that these proteins can activate small GTPases, which regulate many cellular pathways related to nutrient signaling and vesicular trafficking (58). Additional domains found in these proteins include TPR repeats, IG-like domains and HTH/PCI domains which can be involved in interactions with protein substrates for further trafficking (59–61). The SB-containing proteins might interact with SB motifs in ESCRT-I, whereas other domains are likely involved in substrate recognition (Figure 6B). Notably, the distribution of all these families of ESCRT components across Asgard genomes is extremely patchy such that many domain combinations are present in only a few species and rarely in multiple lineages (Figure 6A). Nevertheless, most of the Asgard lineages encompass at least one representative of each of the major ESCRT components, that is, ESCRT-I, -II, -III, -IV.

### Evolution of ESCRT systems

Despite the patchy distribution of the ESCRT systems in archaea, the topology of the Vps4 tree is largely consistent with the species tree (62) (Figures 3A and 7, Supplementary Figure 1, Supplementary File 1), suggesting that these systems are mostly vertically inherited, with multiple lineages-specific losses occurring during evolution (Supplementary Figure 1, Supplementary File 1, Supplementary Table 1). We used the topology of the Vps4 tree as the scaffold for reconstructing an evolutionary scenario for the ESCRT systems based on the parsimony principle (minimal number of events).

**Figure 7.**
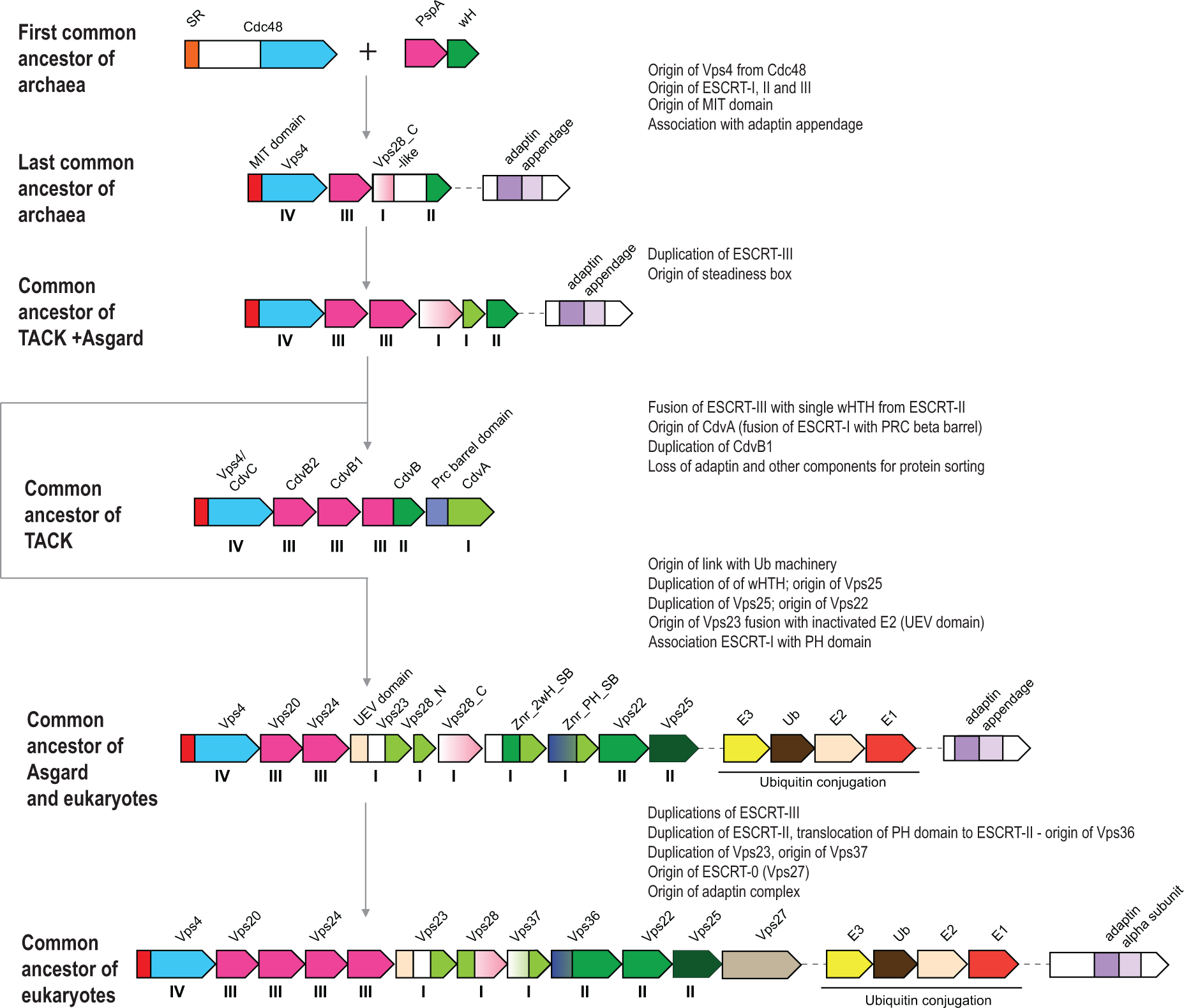
Origin and evolution of ESCRT systems. The inferred, hypothetical organizations of ESCRT systems in ancestral archaeal groups indicated on the left are shown. The inferred key events are described on the right. Designations of the genes and domains are the same as in Figure 3. The Roman numerals under the genes correspond to (putative) components of the four ESCRT complexes. Abbreviations are the same as in Figure 3 except for SR, substrate recognition domain.

ESCRT systems (containing both *vps4* and ESCRT-III genes) are confined to archaea and eukaryotes, and likely evolved in early archaeal ancestors, antedating the last archaeal common ancestor (LACA). As noted above, PspA, the apparent ancestor of ESCRT-III components, can be traced to the LUCA and initially was probably involved in membrane repair (25, 63). Recent analysis has shown that PspA proteins are often encoded next to wH proteins in both bacteria and archaea (63), suggesting a functional link between these two components of future ESCRT systems. In these early archaeal ancestors, Vps4 likely originated from the Cdc48 ATPase, as can be inferred from the presence of a substrate recognition domain (double psi beta-barrel fold), a possible relic of the ancestral state in the Halo_adaptin Vps4-like ATPase, and the overall similarity between Vps4 and Cdc48 (64). One of the two ATPase domains of Cdc48 was lost during this transition. Thus, an early ancestral ESCRT system could consist of Vps4, ESCRT-III and a wH containing protein, the ancestor of ESCRT-II (Figure 6). The wH domain is a key building block found in most ESCRT systems that likely interacts with ESCRT-III (Figure 2 and 4A). Indeed, in many TACK archaea, the wH domain is fused to the C-terminus of ESCRT-III. Subsequently, other components were acquired, resulting in the specialization for different cargo proteins and increasingly sophisticated regulation of the system assembly. One of these components is a coiled-coil domain, which is found in most ESCRT systems and could form a structure analogous to the stalk of ESCRT-I (Figure 1 and 3B). Other ancestral modules could be alpha-helical domains similar to Vps28-C (helical bundle) and the SB box (helical hairpin). These simple structural elements often comprise N- or C-terminal domains in various proteins from archaeal ESCRT systems (Figure 5B, Supplementary Figures 4-7). The exact origin of these elements is difficult to pinpoint, but they might have been derived from ESCRT-III or the MIT domain. Adaptin appendage structural domains were identified in many systems and thus could be functionally linked to early archaeal ESCRT (Figure 3B, 4B and 7).

Thus, the LACA ESCRT system could be similar in complexity to the Meth_adaptin system and probably coordinated protein sorting (Figure 3B). Subsequent diversification of ESCRT yielded specialized systems for phosphorylated protein sorting, in the Main clade and Halo_FHA systems, whereas in TACK and Thermoplasmata, ESCRT underwent reductive evolution (Figure 6). In the TACK ancestor, ESCRT was co-opted for cell division and likely lost most of the components involved in protein sorting. The functions of the reduced ESCRT system in Thermoplasmata remain to be elucidated, but it seems likely to be involved in cell division as well (Figure 5, Supplementary Table 1 and 2).

The ESCRT systems of Asgard archaea are highly complex, but this complexity is achieved mostly by duplications of the basic building blocks described above rather than by acquisition of new components (Figure 7). First, a duplication produced two paralogous wH proteins, which further duplicated giving rise to a eukaryote-like, three-component ESCRT-II complex. Second, the SB box was duplicated and fused to VPS28_C domain or coiled-coil domain, giving rise to the ortholog of eukaryotic Vps28 and Vps23-like protein, respectively. Already at an early stage of Asgard evolution, ESCRT apparently became specialized for ubiquitinated protein sorting, primarily, as a result of the emergence of the UEV domain from the E2 component of the Ub system (65). The UEV domain eventually fused to a Vps23-like protein yielding the ortholog of eukaryotic Vps23. The most complex feature of the Asgard ESCRT-I is the diversity of multidomain proteins containing either an SB or the Vps28_C domain (Figure 6B). Like in Meth_adaptin systems, many of these proteins also contain an adaptin appendage domain. Given that the adaptin appendage domain is a known hub of protein interactions (66, 67), the diversity of these components might provide additional flexibility of interactions with different types of cargo and can be considered functional analogs of ESCRT-0.

The key components of the ESCRT system continued to duplicate and subfunctionalize during the evolution of the major lineages of eukaryotes; distinct ESCRT-0 components emerged, providing an interface for clathrin and clathrin adaptor complexes, the latter evolving from the adaptin appendage domain (Figure 1 and 6).

### Functions of ESCRT systems in archaea

During the recent decade, considerable progress has been reached in understanding the functionality of both FtsZ-based and ESCRT-based molecular machineries in archaea (68). Two independent recent studies reported PRC barrel protein to be essential for cell division in *Haloferax volcanii* (51, 69). It has been shown that two PRC barrel proteins form a tripartite complex with SepF which recruits FtsZ to the divisosome (51, 69). It has been thought previously that Thermoproteales had a unique cell division apparatus because neither FtsZ nor ESCRT system genes were identified in these genomes (24). It has been hypothesized that these organisms used an actin homolog (crenactin hereinafter) along with three other genes (arcadin 1, 2, 3 and 4, which are often encoded in vicinity) for cell division (24, 70). However, experimental analysis of these protein led to inconclusive results (70, 71) so that the role of actin in cell division as well as the functions of arcadin 1, 3 and 4 remain unclear. As indicated above, PRC barrel is an N-terminal domain of CdvA, and recently, a diverged CdvA ortholog has been identified in Thermoproteales (51) (Figure 2 and 5). Apart from the PRC barrel domain in CdvA, several Thermoproteales genomes encode additional PRC barrel proteins (Figure 2, Figure 5B). These proteins are potential additional components of the cell division machinery in this archaeal lineage.

Whereas most of euryarchaea apparently encompass the FtsZ-based division machinery, with all three currently known components (FtsZ, SepF and PRC barrel) conserved, several Thermoplasmata genomes encode only FtsZ but not SepF and PRC, and some others lack all three components (Figure 5A). The latter group includes genomes of three genera, *Picrophilus*, *Acidiplasma* and *Ferroplasma*. *Acidiplasma* and *Ferroplasma* encode a minimal ESCRT system described above along with actin (Figure 5A). Considering that this ESCRT system, similarly to the Cdv system, lack components required for involvement in protein sorting, it might have been co-opted for cell division in these organisms. However, other functions cannot be ruled out because *Thermoplasma volcanium*, *T.acidophilum* and Thermoplasmatales archaeon Gpl (*Cuniculiplasma divulgatum* genomes encoded this system along with the complete FtsZ machinery. The only euryarchaeal genome that lacks all genes implicated in cell division is *Picrophilus torridus* (Figure 5A). The only candidate for involvement in cell division we identified in this genome is a member of Thermoplasmatales-specific uncharacterized arCOG07400 family (WP_011177753.1).

All other complex ESCRT systems are present in organisms that also possess a (mostly) complete FtsZ cell division machinery suggesting that they are not involved in cell division, but rather, in vesicle formation and protein sorting (Supplementary Table 1). In particular, like Thaumarchaea, most Asgard archaea encode FtsZ and SepF (and some PRC barrel proteins) along with ESCRT (Supplementary Table 1). However, unlike in Thaumarchaea that use ESCRT for cell division, Asgard ESCRT systems are likely dedicated to vesicular transport of protein cargo, mostly, ubiquitinated proteins, whereas cell division in Asgard archaea likely relies solely on the FtsZ machinery. However, given that in eukaryotes, the ESCRT system is multifunctional and is involved in cytokinetic abscission (18, 19, 72, 73), the ESCRT system of Asgard archaea also might contribute to cell division.

Many archaea produce vesicles by budding (74), but only in Sulfolobales, the ESCRT system was found to be responsible for the formation of these vesicles suggesting multiple roles of the ESCRT system in these organisms (75, 76). Although many other archaea, especially Thermococcales, in which vesicles were observed (74), do harbor at least some components of the ESCRT machinery, others, such as *Halorubrum lacusprofundi* and *Aciduliprofundum boonei* do not, making it unlikely that ESCRT is solely responsible for vesicle formation in archaea. ESCRT systems in the Main and FHA group systems are associated with S/T kinases, suggesting that protein phosphorylation is key to their functions. The fusion of the FHA domain with the CdvA-like domain suggests that ESCRT system recognize phosphorylated proteins as cargo. In contrast, in the case of Halo_adaptin and Meth_adaptin systems, the cargo is probably bound via protein-protein interactions involving IG-like or PQQ-like domains (Figure 3B). Considering that Halo_adaptin system is the most diverged one and is present in archaea that divide using the FtsZ system, this ESCRT system is likely involved in vesicle formation.

## Concluding remarks

Our analysis of the distribution, domain architectures and genomic neighborhoods of ESCRT components in archaea revealed striking evolutionary flexibility and modularity, most conspicuously, in Asgardarchaeota. Although undoubtedly subject to amendment by comparative analysis of an even greater diversity of archaeal genomes yet to be sequenced, the available data is sufficient to develop a plausible scenario of the origin and evolution of the ESCRT machinery.

The ESCRT system involved in protein sorting is an early archaeal innovation. Only ESCRT-III antedates archaea, deriving from a PspA-like protein that was probably involved in membrane repair in the LUCA (25, 63). We can now place ESCRT-II and adaptin appendage domains into the LACA and hypothesize that the LACA also possessed analogs of Vps28-C (alpha helical bundle). The exact interactions between the components in different archaeal ESCRT systems remain to be elucidated. We identified strong contextual links between ESCRT and such domains as GOLD and SPFH, suggesting that building blocks for these eukaryotic signature systems originated early in archaeal evolution. The ESCRT machinery of the LACA was likely involved primarily in protein sorting via endocytosis. The subsequent evolution of ESCRT took divergent paths in different major lineages of archaea. In TACK and Thermoplasmata, the ancestral ESCRT system seems to have evolved towards secondary simplification, whereby only the core components of the system were retained whereas all putative ESCRT-0 analogs were lost. In these groups of archaea, ESCRT was co-opted for cell division and, at least in Sulfolobales, for extracellular vesicle biogenesis (33). Notably, this reduction of the ESCRT system and its repurposing for cell division occurred on at least two independent occasions in the evolution of archaea. Conversely, in Asgard archaea, ESCRT evolved towards complexification, whereby ESCRT complexes subfunctionalized to mediate membrane remodeling through interaction with the actin-based cytoskeleton(77). Furthermore, the link between ESCRT and the ubiquitin network, presently considered a signature of the eukaryotic cell biology, likely evolved in Asgard archaea (22, 27, 28). Asgard archaea also possess multiple ESCRT components that apparently have not been inherited by eukaryotes (Figure 2A and 6B). The complex relationship between the archaeal and eukaryotic ESCRT machineries described here contrasts the more direct correspondence between the components of several other conserved macromolecular complexes, such as the proteasome (78) or the exosome (79). This distinction likely reflects pronounced differences between membrane remodeling and protein sorting processes in Asgard archaea and eukaryotes, despite the conservation of the core machinery.

Multiple recent phylogenetic analyses based on a variety of marker genes converge on the conclusion that eukaryotes are either the sister clade of the Heimdall supergroup, which includes Heimdallarchaeia, Kariarchaeia, Gerdarchaeia and Hodarchaeia lineages, or the sister clade of Hodarchaeia within the Heimdall supergroup (27–30). However, our phylogenetic analysis of Vps4 and previous analyses of ESCRT III (25, 63) and ESCRT II components (22, 27) suggest affinity between eukaryotes and Asgardarchaeota as a group rather than with the Heimdall supergroup or Hodarchaeia. Furthermore, we did not detect any synapomorphies (derived shared characters such as common domain architectures) between either the Heimdall supergroup or any other Asgard lineages and eukaryotes. On the contrary, there are several distinct features in the organization of the core ESCRT machinery in Asgards as discussed above. So far, the conclusion of origin of eukaryotes within the Heimdall supergroup or from a common ancestor with Hodarchaeia is solely based on phylogenetic analysis of (nearly) universal genes, but no systematic search for synapomorphies shared between eukaryotes and different Asgard lineages has been reported. Considering the essential role of ESCRT in both Asgard archaea and eukaryotes, analysis of ESCRT components suggests extra caution regarding the inference of the exact nature of the evolutionary relationship between Asgardarchaeota and eukaryotes. A comprehensive search for synapomorphies to complement the phylogenetic analyses should help clarify these relationships and shed more light on the eukaryotic ancestry.

## Methods

### Orthologous gene datasets

The arCOGs and asCOGs databases were used as the comparative genomic framework (27, 34, 80). The arCOGs are annotated clusters of orthologous genes from 524 archaeal genomes covering all major archaeal lineages whereas asCOGs are annotated clusters of orthologous genes from 78 genomes of Asgardarchaeota covering all major Asgard lineages.

### Genome and protein sequence analysis

PSI-BLAST (81) was used to identify homologs of all known ESCRT proteins and homologs of all proteins found in their neighborhoods in the arCOGs database. Given that the sequences of many ESCRT proteins are compositionally biased, the searches were run with variable parameter sets and different queries. Typically, PSI-BLAST was run for 5 iterations or until convergence, with inclusion threshold E-value=0.0001 and compositional adjustments statistics turn off. Low scoring hits were manually examined and verified using HHpred (82). HHpred search was run on the web against PDB (83), CDD v3.16 (84) and PFAM_A v36 (85) derived profile libraries imbedded in the HHpred toolkit with otherwise default parameters. All HHpred hits were visually examined, and the highest-scoring relevant hit is reported for newly identified domains (Supplementary Table 3). The Marcoil program (86) was used to predict coiled-coil regions and TMHMM (87) was used to predict transmembrane segments. Domain boundaries were determined based on the combination of evidence including examination of HHpred hits and comparison of output alignments with the respective structures from the PDB database (when available), examination of AF2 models and transmembrane and coiled-coil regions predicted using TMHMM and Marcoil, respectively. Based on this analysis, 11 arCOGs and one asCOG (cog.002483) were corrected, and 6 new arCOGs (arCOG15281-6) were created.

In order to increase the diversity of archaeal ESCRT groups identified here (Meth_adaptin and Halo_adaptin), the NCBI NR database was searched for closely related homologs in database using Vps4 ATPase sequences from respective genomes as queries. Respective genomic loci were examined to ascertain that they encoded all or most of the identified ESCRT components from the respective groups. These loci and the respective protein accessions are listed in Supplementary Tables 2 and 3.

### Phylogenetic analysis

The Vps4 protein sequences were the clustered using Mmseqs2 (88) with a 90% protein identity threshold, and a representative of each cluster was included in the sequence set for phylogenetic tree reconstruction. Muscle5 (89) with default parameters was used to construct the multiple alignment of the Vps4 family. For phylogenetic analysis, several poorly aligned sequences or fragments were discarded. Columns in the multiple alignment were filtered for homogeneity value (90) of 0.05 or greater and a gap fraction less than 0.667. This filtered alignment was used as the input for the IQ-tree (91) to construct maximum likelihood phylogenetic tree with the LG+F+R10 evolutionary model (Supplementary File 1, Supplementary Table 1). The same program was used to calculate support values. To maximize the number of phylogenetically informative positions for the set containing only Vps4 sequences from archaea and eukaryotes, with the position of the root determined by building a second tree which contained only archaeal Vps4 sequences from the arCOGs and several selected Cdc48 family ATPases (C-terminal domain), the closest homolog of Vps4, as an outgroup (Supplementary Figure 1, Supplementary File 2, Supplementary Table 1). The alignment for this tree was filtered as described above and the tree was reconstructed using IQ-tree with the LG+R6 best fitted evolutionary model. The Vps4 tree shown on the Figure 3A was then rooted according to the root position found in the tree that included Cdc48 as an outgroup (Supplementary Figure 1).

### Protein structure prediction and analysis

For protein structure predictions, all relevant sequences were extracted from the respective gene neighborhoods (Supplementary Table 2) and queried against a combination of databases (UniRef30-2022/05 (92), pdb70-2022/03/01 (93) and mgnify-2022/05 (94)) using HHblits (93). The resulting MSAs were used as input for Alphafold2.3.1 (31) using the monomer model and generating 5 model structures per alignment. For structure visualizations, the overall highest pLDDT scoring model was used. USCF ChimeraX was used for all structural analysis and visualization (95). Structure comparisons were performed using DALI (96).

## Data Availability

The data used in this work are available through public databases or the Supplementary Material. Additional Supplementary files (Supplementary File 1, 2 and Asgard sequences corresponding to Supplementary Table 2) are available via ftp: https://ftp.ncbi.nih.gov/pub/makarova/Supplement/ESCRT/

## Author contributions

K.S.M and E.V.K initiated the project; K.S.M. collected the data; K.S.M., V.T., Y.I.W, Z.L., Y.L., S.Z., M.K., M.L. and E.V.K. analyzed the data; K.S.M., V.T. and E.V.K. wrote the manuscript that was read, edited, and approved by all authors.

## Acknowledgments

This work was supported by the National Natural Science Foundation of China (Grant Nos. 32225003,32000002,92251306, 31970105), Guangdong Basic and Applied Basic Research Foundation (Grant No. 2023A1515011309), the Innovation Team Project of Universities in Guangdong Province (No. 2020KCXTD023), and the Shenzhen Science and Technology Program (Grant no. JCYJ20200109105010363). K.S.M., V.T., Y.I.W. and E.V.K. are supported by intramural funds of the US Department of Health and Human Services (National Institutes of Health, National Library of Medicine).

## Supplementary Material

**Supplementary Table 1: ESCRT and cell division system components in archaea. The** first worksheet contains information on the presence and absence, and the number of paralogs for each gene in these systems is provided for each of 524 archaeal genomes. The second worksheet contains the list of all components of ESCRT systems (protein accession, taxonomy information and other metadata is provided. The third and fourth worksheets contain, information for the sequences included in the trees in the Supplementary Figure 1 and Figure 3A, respectively.

**Supplementary Table 2: Annotated neighborhoods of ESCRT genes.** Five genes upstream and downstream of the respective ESCRT gene for genomes in arCOGs and asCOGs are shown.

**Supplementary Table 3: Evidence of sequence similarity for predicted components of archaeal ESCRT systems.**

**Supplementary Figure 1: Phylogenetic tree of archaeal Vps4 tree rooted using Cdc48 as an outgroup.** The maximum likelihood phylogenetic tree was built for 176 archaeal Vps4 sequences using IQ-tree program with the LG+R6 best fitted evolutionary model (Supplementary Table 2, Supplementary File 2). The same program was used to calculate support values. The tree was rooted using Cdc48 C-terminal ATPase domain from selected archaea (gray). The major clades corresponding to distinct ESCRT systems is denoted on the right. The species were colored according to their taxonomy which is also indicated on the right with the same color. A few species that do not belong to these major archaeal taxa remain black.

**Supplementary Figure 2: Structural models for the main clade of ESCRT systems.** The gallery of structure predictions obtained for the Main clade (Figure 3A) using the gene neighborhood from *Palaeococcus pacificus* as representatives (Supplementary table 3) is shown. Gene neighborhood organization is shown on top. Proteins are colored by common structural domains found in ESCRT gene neighborhoods. Named proteins are assigned by sequence or structural similarity. NCBI accessions are shown for all predictions. Unstructured termini and long linkers are hidden. Abbreviation are as per previous legends, with the addition of HD, Helical Domain and TMH, Transmembrane helix.

**Supplementary Figure 3: Multiple Alignment of arCOG08177 sequences**

Cas2 and TBP domains are indicated above the alignment. The alignment was colored using http://www.bioinformatics.org/sms2/color_align_cons.html tool with default parameters for amino acid grouping and consensus 80%. The alignment with two previously studied Cas2 (104, 105) is based on HHpred results. The positions corresponding to catalytic residues of Cas2 are highlighted in red. A few amino acids from the C-terminus of several sequences are omitted.

**Supplementary Figure 4: Alignment and structural models for CdvA helical hairpin and Steadiness box (SB).**

Manual alignment of SB-like alpha helix hairpin is based on the respective HHpred alignments with profile PF18822.4 corresponding to CdvA. Protein accession, locus tag, gene name or PDB ID (for eukaryotes) are indicated on the left. Major taxonomic lineages are indicated for each sequence on the right. Details for the respective HHpred searches are provided in the Supplementary Table 4. Alpha helices are denoted by “@” above the alignment. Alignment coloring is based on default amino acid groups implemented in http://www.bioinformatics.org/sms2/color_align_cons.html tool with 50 % consensus. Structural models of selected proteins containing SB or CdvA are shown. The SB-like hairpin is shown in blue. ESCRT III protein, which also has a distinctive alpha helix hairpin is shown for comparison.

**Supplementary Figure 5: Structural models for the Halo_FHA clade**

The gallery of structure predictions obtained for the Halo_FHA clade (Figure 3A) using the gene neighborhood from *Natronolimnobius baerhuensis* as representatives (Supplementary Table 3) is shown. The gene neighborhood organization is shown on top. Proteins are colored by common structural domains found in the ESCRT gene neighborhoods. Protein names are assigned by sequence or structural similarity. Unstructured termini and long linkers are hidden. Abbreviation are as per previous legends with addition of: HD, Helical Domain; TMH, Transmembrane helix

**Supplementary Figure 6: Structural models for the Meth_adaptin clade**

The gallery of structure predictions obtained for the Meth_adaptin clade (Figure 3A) using the gene neighborhood from *Methanosalsum zhilinae* as representatives is shown (Supplementary table 3). The gene neighborhood organization is shown on top. Proteins are colored by common structural domains found in the ESCRT gene neighborhoods. Protein names are assigned by sequence or structural similarity. Unstructured termini and long linkers are hidden. Abbreviation are as per previous legends with addition of: HD, Helical Domain; TMH, Transmembrane helix.

**Supplementary Figure 7: Structural models for the Halo_adaptin clade**

The gallery of structure predictions obtained for the Halo_adaptin clade (Figure 3A) using the gene neighborhood from *Haloquadratum walsbyi* as representatives is shown (Supplementary Table 3). The gene neighborhood organization is shown on top. Proteins are colored by common structural domains found in the ESCRT gene neighborhoods. Proteins names are assigned by sequence or structural similarity. Unstructured termini and long linkers hidden. Abbreviation are as per previous legends with the addition of HD, Helical Domain; TMH, Transmembrane helix.

